# Serum Amyloid P inhibits single stranded RNA-induced lung inflammation, lung damage, and cytokine storm in mice

**DOI:** 10.1101/2020.08.26.269183

**Authors:** Tejas R. Karhadkar, Darrell Pilling, Richard H. Gomer

## Abstract

SARS-CoV-2 is a single stranded RNA (ssRNA) virus and contains GU-rich sequences distributed abundantly in the genome. In COVID-19, the infection and immune hyperactivation causes accumulation of inflammatory immune cells, blood clots, and protein aggregates in lung fluid, increased lung alveolar wall thickness, and upregulation of serum cytokine levels. A serum protein called serum amyloid P (SAP) has a calming effect on the innate immune system and shows efficacy as a therapeutic for fibrosis in animal models and clinical trials. In this report, we show that aspiration of the GU-rich ssRNA oligonucleotide ORN06 into mouse lungs induces all of the above COVID-19-like symptoms. Men tend to have more severe COVID-19 symptoms than women, and in the aspirated ORN06 model, male mice tended to have more severe symptoms than female mice. Intraperitoneal injections of SAP starting from day 1 post ORN06 aspiration attenuated the ORN06-induced increase in the number of inflammatory cells and formation of clot-like aggregates in the mouse lung fluid, reduced ORN06-increased alveolar wall thickness and accumulation of exudates in the alveolar airspace, and attenuated an ORN06-induced upregulation of the inflammatory cytokines IL-1β, IL-6, IL-12p70, IL-23, and IL-27 in serum. Together, these results suggest that aspiration of ORN06 is a simple model for both COVID-19 as well as cytokine storm in general, and that SAP is a potential therapeutic for diseases with COVID-19-like symptoms as well as diseases that generate a cytokine storm.

## Introduction

Coronavirus disease 2019 (COVID-19) is generally caused by inhalation of the severe acute respiratory syndrome coronavirus 2 (SARS□CoV□2), a single-stranded RNA (ssRNA) virus related to severe acute respiratory syndrome coronavirus (SARS-CoV) and Middle East respiratory syndrome-related coronavirus (MERS-CoV) (1). Human patients and murine models of SARS-CoV, MERS, and SARS-CoV2 indicate that severe disease may be triggered by a cytokine storm, in which, as in other cytokine storm associated diseases, an over exuberant production of cytokines leads to the accumulation of immune cells in sensitive organs such as the lungs (2, 3). Although there are similar numbers of confirmed SARS-CoV2 cases between men and women, severe COVID-19 disease, as measured by hospitalization, admission to intensive care units, and rates of mortality, are 1.5 to 2 times higher in men than women (4-7). These data are independent of other known factors, such as smoking, diabetes, obesity, heart disease, chronic kidney disease, or chronic lung disease (7, 8).

Lung biopsy and autopsy findings indicate that COVID-19 is associated with an accumulation of cell-free protein-rich exudate in the alveoli, alveolar wall thickening, and inflammation (accumulation of immune cells) (9-15). COVID-19 patients are also at risk of disseminated intravascular coagulation and deposition of fibrin clots in the lungs (9, 10, 14-16). In both animal models and COVID-19 patients, there is a consistent accumulation of macrophages in the lungs, whereas increased numbers of lung neutrophils appear to be associated with comorbidities such as pneumonia (14, 17, 18).

Serum amyloid P (SAP; PTX2) is a member of the pentraxin family of proteins that includes C-reactive protein (CRP; PTX1) and pentraxin-3 (PTX3). SAP is made by hepatocytes and secreted into the blood (19, 20). SAP binds to a variety of molecules including DNA and apoptotic debris, bacterial polysaccharides, amyloid deposits, and bacterial and viral proteins (21-23). Phagocytic cells such as monocytes and macrophages then bind the SAP and engulf the debris or other material the pentraxin has bound (24). SAP binds to Fc receptors (receptors for IgG) as well as the C-type lectin receptor DC-SIGN/CD209 on monocytes and macrophages (24-28), and inhibits the formation of pro-inflammatory and pro-fibrotic macrophages, and promotes the formation of immuno-regulatory macrophages (29-40). In addition to promoting the formation of immuno-regulatory macrophages, SAP induces macrophages to secrete the anti-inflammatory cytokine IL-10 (34, 35, 41). SAP decreases neutrophil binding to extracellular matrix components (42-44), and in a mouse model of lung inflammation, SAP injections starting 24 hours after insult reduced the number of neutrophils in the lungs (44). In animal models and human clinical trials, SAP inhibits fibrosis (30, 35, 45-47). SAP is also effective as an inhibitor of influenza A virus infections (48-50).

Cells can sense the presence of the SARS□CoV□2 ssRNA virus using the ssRNA-sensitive intracellular toll-like receptors TLR7 and TLR8 (51, 52). Adding GU-rich ssRNAs to human peripheral blood immune cells induces proinflammatory cytokine production, and tail vein injections of GU-rich ssRNAs in mice lead to pulmonary edema, accumulation of inflammatory cells, and alveolar hemorrhage/damage in the lung (51, 53).

In this report, we show that oropharyngeal aspiration of ORN06 induces a cytokine storm and lung damage in mice with remarkable similarities to the cytokine storm and lung damage seen in COVID-19 patients, including more severe effects in male mice than female mice. We find that SAP strongly attenuates the ORN06 effects, suggesting that SAP could be a potential therapeutic for severe COVID-19 and possibly cytokine storms in general.

## Materials and Methods

### Preparation of SAP

Purified human SAP (30C-CP1104LY) was purchased from Fitzgerald Industries (Acton, MA). Before use, the SAP was buffer exchanged into 20 mM sodium phosphate, pH 7.4 to remove sodium azide and EDTA present in the commercial preparation. The SAP was diluted 1:1 with 20 mM sodium phosphate, pH 7.4, and added to Amicon Ultra 0.5ML 10 kDa cutoff centrifugal filter units (Millipore-Sigma, Burlington, MA) and centrifuged at 10,000 x g for 10 minutes. The SAP in the upper reservoir was then resuspended with 0.4 ml 20 mM phosphate buffer and centrifuged again. This process was repeated four times to exchange the buffer to 20 mM sodium phosphate. Buffer-exchanged preparations were then checked for protein concentration by OD 260/280/320 using a Take3 micro-volume plate with a SynergyMX plate reader (BioTek, Winooski, VT, USA) as previously described (54). The SAP, at 2.5-4 mg/ ml, was stored in aliquots at 4 °C and used within 14 days. SDS-PAGE of 1 µg of this SAP showed a single band at the expected molecular mass of 26 kDa.

### Mouse model of lung damage and cytokine storm

5-week old 20 g male and female C57BL/6 mice (Jackson Laboratories, Bar Harbor, ME) were given an oropharyngeal aspiration of 1 mg/kg (or less where indicated) ORN06/LyoVec (a ssRNA oligonucleotide with 6 UGU repeats complexed with a lipid which protects the ssRNA from degradation; henceforth referred to as ORN06) (InvivoGen, San Diego, CA) in 50 µl phosphate-buffered saline (PBS), or an oropharyngeal aspiration of an equal volume of PBS as a control, following (55, 56). At 24 and 48 hours after ORN06 insult, the control mice and some of the ORN06-treated mice were given intraperitoneal injections of 100 µl PBS, or SAP diluted to a dose of 2.5 mg/kg in 100 µl PBS. At day 3, mice were euthanized by CO_2_ inhalation, and bronchoalveolar lavage fluid (BALF) was obtained as previously described (55, 57). The experiment was performed in accordance with the recommendations in the Guide for the Care and use of Laboratory Animals of the National Institutes of Health. The Texas A&M University Animal Use and Care Committee approved the protocol.

### Staining of bronchoalveolar lavage fluid (BALF) cells

BALF cells were counted and processed to prepare cell spots as described previously (58, 59). After air drying for 48 hours at room temperature, some of the prepared BALF cell spots were fixed and treated with Wright-Giemsa stain (Polysciences, Inc., Warrington, PA) following the manufacturer’s instructions. At least 150 cells per mouse were examined and quantified for cell type following the manufacturer’s instructions using a CME microscope (Leica, Buffalo Grove, IL) with a 40x objective, and selected areas were imaged with a 100x oil immersion objective on an Eclipse Ti2 microscope (Nikon, Melville, NY) with a 10 megapixel color CCD camera (Amscope, Irvine, CA).

Immunochemistry on BALF cell spots and counts of cells was performed as described previously (57, 60, 61). Slides were incubated with monoclonal antibodies overnight at 4°C using anti-CD3 (100202, clone 17A2, BioLegend, San Diego, CA) to detect T-cells, anti-CD11b (101202, clone M1/70, BioLegend) to detect blood and inflammatory macrophages, anti-CD11c (M100-3, clone 223H7, MBL International, Woburn, MA) to detect alveolar macrophages and dendritic cells, anti-CD45 (103102, clone 30-F11, BioLegend) for total leukocytes, anti-Ly6G (127602, clone 1A8, BioLegend) to detect neutrophils, with isotype-matched irrelevant rat antibodies as controls, diluted to 5 µg/ml in PBS containing 2% (w:v) Fraction V BSA bovine serum albumin (PBS-BSA) (VWR, Radnor, PA). Slides were then washed in PBS, with 6 changes of PBS over 30 minutes, and then incubated with 1 µg/ml biotin-conjugated F(ab’)2-donkey anti-rat antibodies (Novus Biologicals, Littleton, CO) in PBS-BSA for 30 minutes at room temperature. Slides were washed as above and then incubated with 1:1000 streptavidin-conjugated alkaline phosphatase (Vector Laboratories, Burlingame, CA) in PBS-BSA for 30 minutes and washed as above. Staining was developed with the Vector Red Alkaline Phosphatase Kit (Vector Laboratories) for 4 minutes at room temperature following the manufacturer’s directions. Cells were counterstained with Gill’s hematoxylin #3 (Sigma-Aldrich, St. Louis, MO), air dried overnight, and then coverslip (VWR) mounted in Permount mounting medium (17986-01, Electron Microscopy Sciences, Hatfield, PA). Using a 40x objective, at least 100 cells from each stained BALF spot were examined and the percent positive cells was recorded.

### Lung histology

After collecting BALF, the lungs from the mice were harvested and inflated with Surgipath frozen section compound (#3801480, Leica, Buffalo Grove, IL), frozen on dry ice, and stored at −80 °C. 10 µm cryosections of lungs were placed on Superfrost Plus glass slides (VWR) and were air dried for 48 hours. The slides were fixed in acetone for 20 minutes, and then air dried. Slides were incubated for 5 minutes in water at room temperature and then stained for two minutes at room temperature with Gill’s hematoxylin No. 3 solution (Sigma-Aldrich, St. Louis, MO) diluted 1:3 with water and then rinsed under tap water for 1 minute at room temperature. The slides were then incubated with 0.1% eosin for 1 minute and then washed by dipping in water for 10 seconds. The stained slides were allowed to air dry for 2 hours; cleared with xylenes (VWR); and mounted with Permount mounting medium (Electron Microscopy Sciences). After drying the mounted slides for 48 hours, images were captured using a 40x objective on a Nikon Eclipse Ti2 microscope. Images of a 10 µm divisions calibration slide (#MA663, Swift Microscope World, Carlsbad, CA) were used for size bars. Three fields of view were chosen randomly for imaging. Image quantification was done using Image J (NIH, Bethesda, MD). Using the Set Scale function from Image J on a 1 mm scale bar image, all the images were calibrated to the same scale by applying the settings globally. Using the ‘Straight’ command, the length of the alveolar wall on the image was measured and was recorded with the ‘Measure’ command. Eight measurements of the alveolar wall thickness were recorded for each image.

To assess exudates in the alveoli, for each mouse, for the stained cryosections described above, three randomly chosen fields of view were photographed with a 20x objective, and both the number of exudate-containing alveoli and exudate-free alveoli in each image were counted. The percent of exudate-containing alveoli was then calculated. Using the Threshold function and the Measure Tool in NIH ImageJ, the percent area of each 20x objective image of lung cryosections described above containing tissue (not counting pale pink exudates) was measured, and this percentage was subtracted from 100% to obtain the percent area of the image occupied by the alveolar airspace. The threshold was then adjusted to obtain the percent area of the image occupied by tissue and exudate. The percent area of the image occupied by exudate was then calculated by subtracting the percent area of the image occupied by tissue from the percent area of the image occupied by tissue and exudate. The percentage of the exudate-occupied airspace area was then calculated by taking the ratio of (percent area of the image occupied by exudate)/(percent area of the image occupied by the alveolar airspace), and converting the fraction to a percentage.

Lung sections were also assessed for the presence of immune cells using anti-CD11b, anti-CD11c, anti-CD45, anti-Ly6G, or anti-Mac2 (clone M3/38; BioLegend) antibodies with isotype-matched irrelevant rat antibodies as controls. Lung sections were first incubated with PBS containing 4% BSA (PBS-BSA) for 60 minutes to block non-specific binding, and the endogenous biotin was blocked by the addition of streptavidin and biotin solutions, following the manufacturers’ instructions (Streptavidin/Biotin Blocking Kit, Vector Laboratories, Burlingame, CA). Sections were then stained as described above. Positively stained cells were counted from randomly selected fields, and presented as the number of positive cells per mm^2^.

### Serum cytokines

After mice were euthanized, blood was collected from the abdominal aorta and chilled on ice. After 30 minutes, the blood was clarified by centrifugation at 10,000 × g for 5 minutes at 4 °C to isolate serum, which was then stored at −80 °C (62-64). Sera were assayed for cytokines using a 13-plex LEGENDplex Mouse Inflammation Panel kit (Biolegend, San Diego, CA) following the manufacturer’s instructions using an Accuri C6 flow cytometer (BD Bioscience, Franklin Lakes, NJ).

### Statistics

Statistical analyses with t tests or 1-way ANOVA with the indicated post-test were done using GraphPad Prism 7 (GraphPad, San Diego, CA). Significance was defined as p < 0.05.

## Results

### Aspirated ORN06 increases cells in mouse BALF, and SAP injections counteract this

To determine if aspiration of ORN06 can induce lung inflammation and damage, and whether SAP injections might affect this, mice were treated with an oropharyngeal aspiration of ORN06, and then at 24 and 48 hours, mice were treated with intraperitoneal injections of SAP or buffer. Mice were then euthanized at 72 hours. Compared to control (PBS aspirated and then PBS injected), for male and female mice, ORN06 (1 mg/ kg ORN06 aspirated and then PBS injected) caused a large increase in the number of cells recovered from the BALF, and SAP ameliorated the ORN06-induced increase (Fig 1A and S1 Fig). Preliminary experiments with 0.01 and 0.1 mg/ kg ORN06 showed 2.5 × 10^4^ and 3.3 × 10^4^ cells respectively in the BALF for 1 male mouse at each dose. Assessing cell type by Wright-Giemsa staining, for male and female mice, compared to control, ORN06 increased the number of monocyte/macrophages, and SAP treatments reversed these ORN06-induced increases (Fig 1B and S1 Fig). Similar results were observed for lymphocytes, with the exception that control males had more BALF lymphocytes that control females, and the ORN06 treated males showed a wide range of BALF lymphocyte counts and this was thus not statistically different from control (Fig 1C and S1 Fig). The ORN06 treatment did not significantly increase the number of neutrophils in the BALF from male or female mice (Fig 1D and S1 Fig). Compared to ORN06 alone, SAP injections in ORN06-treated mice increased body weight at day 3, with a significant effect in female but not male mice (S2 Fig). For male and female mice, the ORN06 treatment with or without SAP caused no significant difference in liver, heart, kidneys, spleen, white fat, or brown fat weights as a percentage of body weight at day 3 (S3 Fig).

**Fig 1.**
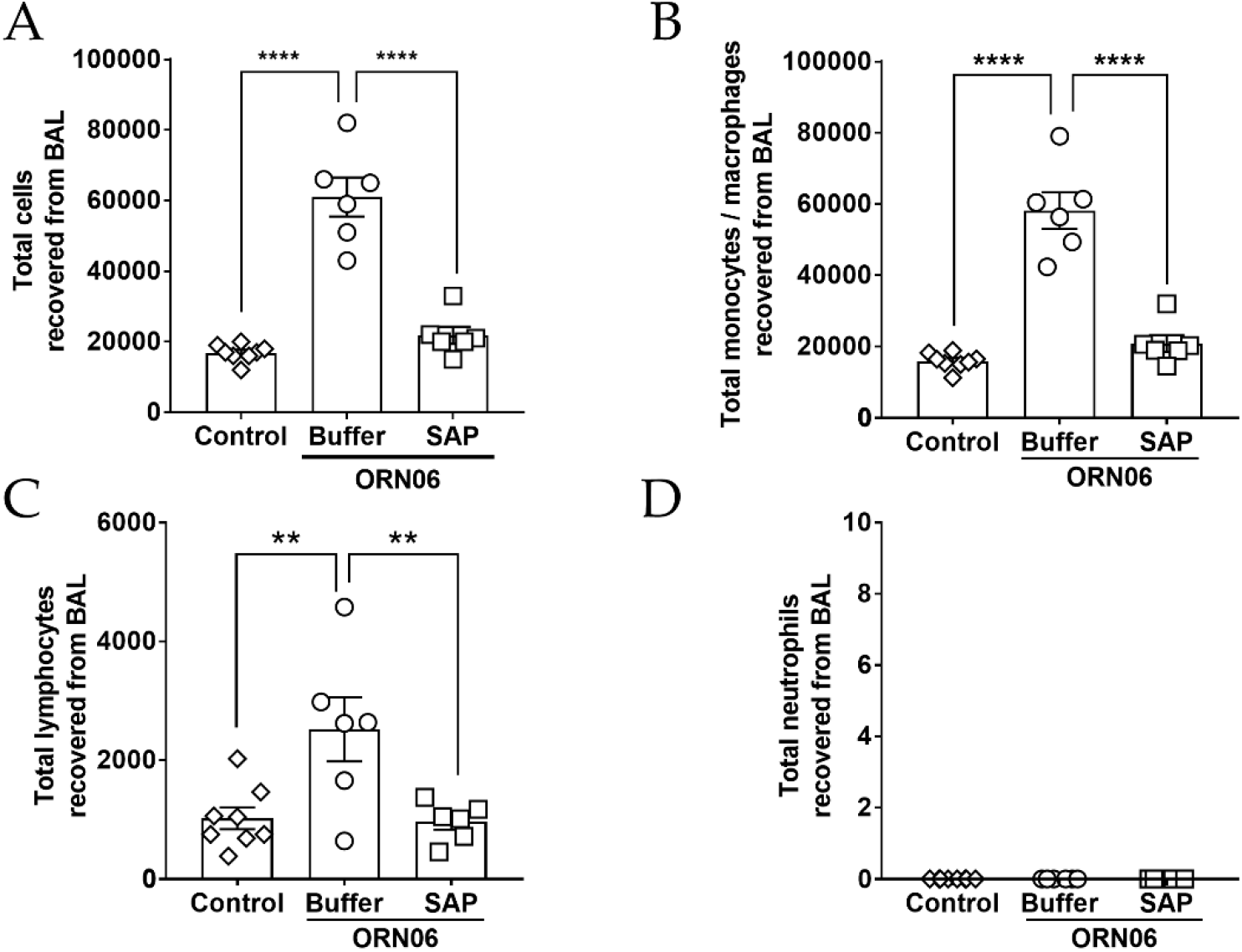
SAP attenuates ORN06-induced upregulation of total cells and immune cell types in mouse BAL. **(A)** The total number of cells collected from the BALF at day 3 post ORN06 aspiration. Cell spots of BALF at day 3 post ORN06 aspiration were stained with Wright-Giemsa stain and the percentage of **(B)** monocytes/macrophages, **(C)** lymphocytes, and **(D)** neutrophils was determined by examining at least 150 cells per mouse BAL sample. Values are mean ± SEM, n = 6 mice (3 males and 3 females), except control group where n = 8 (3 males and 5 females). ** p < 0.01 and **** p < 0.0001 (1-way ANOVA, Bonferroni’s test). For A-D, there was no significant difference between the control group and the SAP-treated group.

Immunohistochemistry on BALF cell spots indicated that compared to control, ORN06 increased the number of CD11c-positive macrophages and/or dendritic cells, and increased the number of CD45-positive leukocytes for both male and female mice, and these effects were reversed with SAP injections (Fig 2 and S4 Fig). Male control mice had more CD11b-positive blood and inflammatory macrophages in the BALF than female control mice (S4 Fig). Compared to control, ORN06 increased the number of CD11b positive cells in the BALF from female mice but not male mice, and SAP treatments reversed the ORN06-induced increase in the female mice (S4 Fig). The ORN06 treatment did not significantly increase the number of CD3-positive lymphocytes or Ly6G-positive neutrophils in the BALF from male or female mice (Fig 2 and S4 Fig). Together, these results indicate that inhaled ORN06 causes an increase in monocytes/macrophages and CD-3 negative lymphocytes in the lung airspace, and that SAP injections can reverse this.

**Fig 2.**
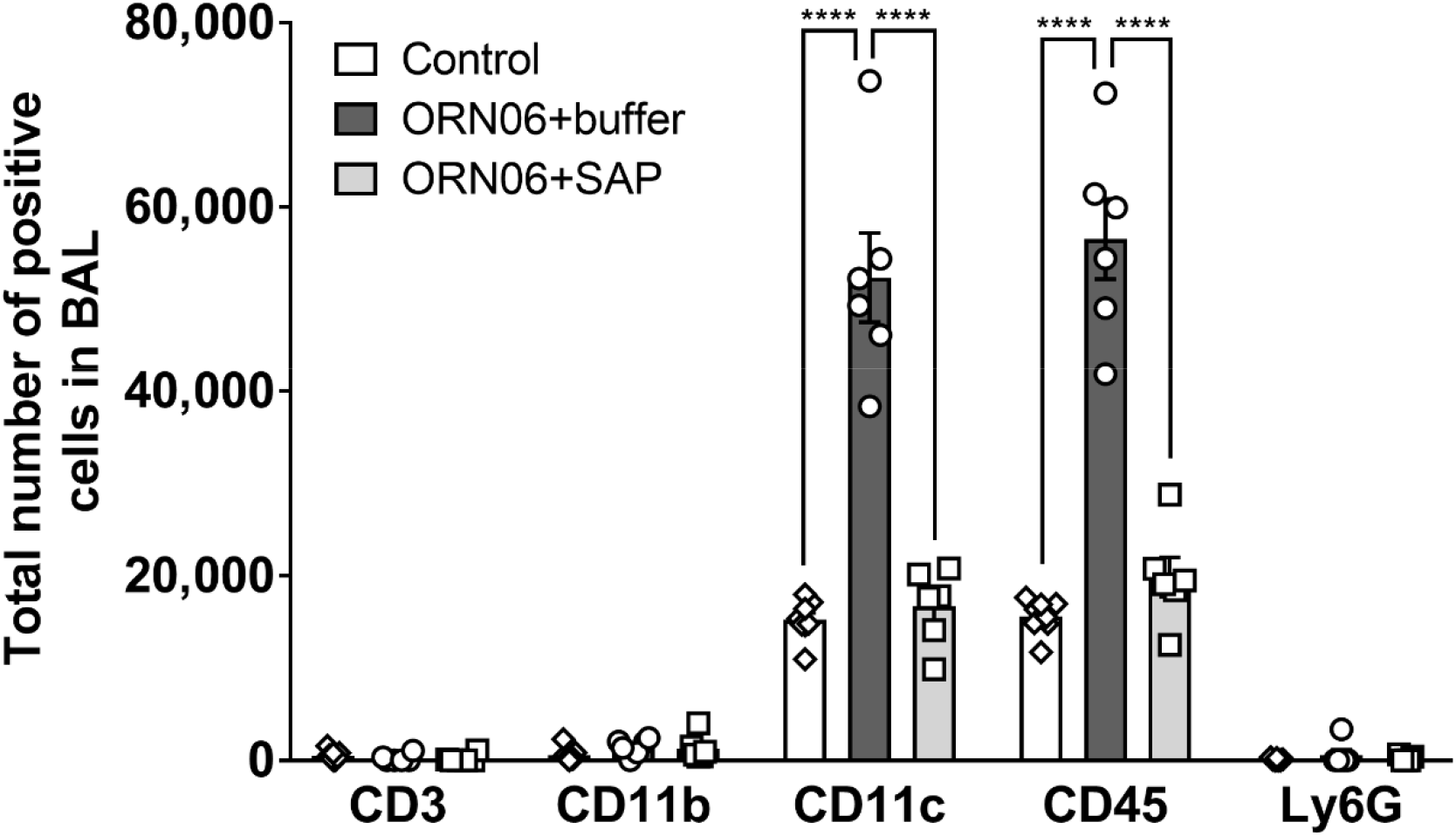
SAP attenuates ORN06-induced upregulation of inflammatory cells in mouse BAL. Cell spots of BALF at day 3 were stained for the indicated markers. The percentage of cells stained was determined in 5 randomly chosen fields of 100–150 cells, and the percentage was multiplied by the total number of BALF cells for that mouse to obtain the total number of BALF cells staining for the marker. Values are mean ± SEM, n = 6 mice (3 males and 3 females), except control group where n = 8 (3 males and 5 females). **** = p < 0.001 (1-way ANOVA, Dunnett’s test). For A-E, there was no significant difference between the control group and the SAP-treated group.

### Aspirated ORN06 increases the local density of macrophages remaining in mouse lungs after BAL, and SAP injections counteract this

After BAL, lungs were inflated and cryosections were immunostained to determine if ORN06 aspiration affects the local density of selected immune cells. Compared to control, ORN06 increased the density of CD11b positive blood and inflammatory macrophages and CD45 positive leukocytes, and SAP reversed this, for male and female mice (Fig 3, S5 Fig, and S6 Fig). The ORN06 treatment with or without SAP caused a greater local density of CD11b positive cells in male compared to female mice, and ORN06 alone caused a greater number of CD45 positive cells in male mice (S6 Fig). Compared to control, ORN06 increased the density of CD11c positive alveolar macrophages and dendritic cells, and SAP reversed this, with significant effects for female but not male mice (Fig 3 and S6 Fig). ORN06 with or without SAP had no significant effect for male or female mice on the local density of Mac2/Gal3 positive alveolar macrophages (65) or Ly6G positive neutrophils (Fig 3 and S6 Fig).

**Fig 3.**
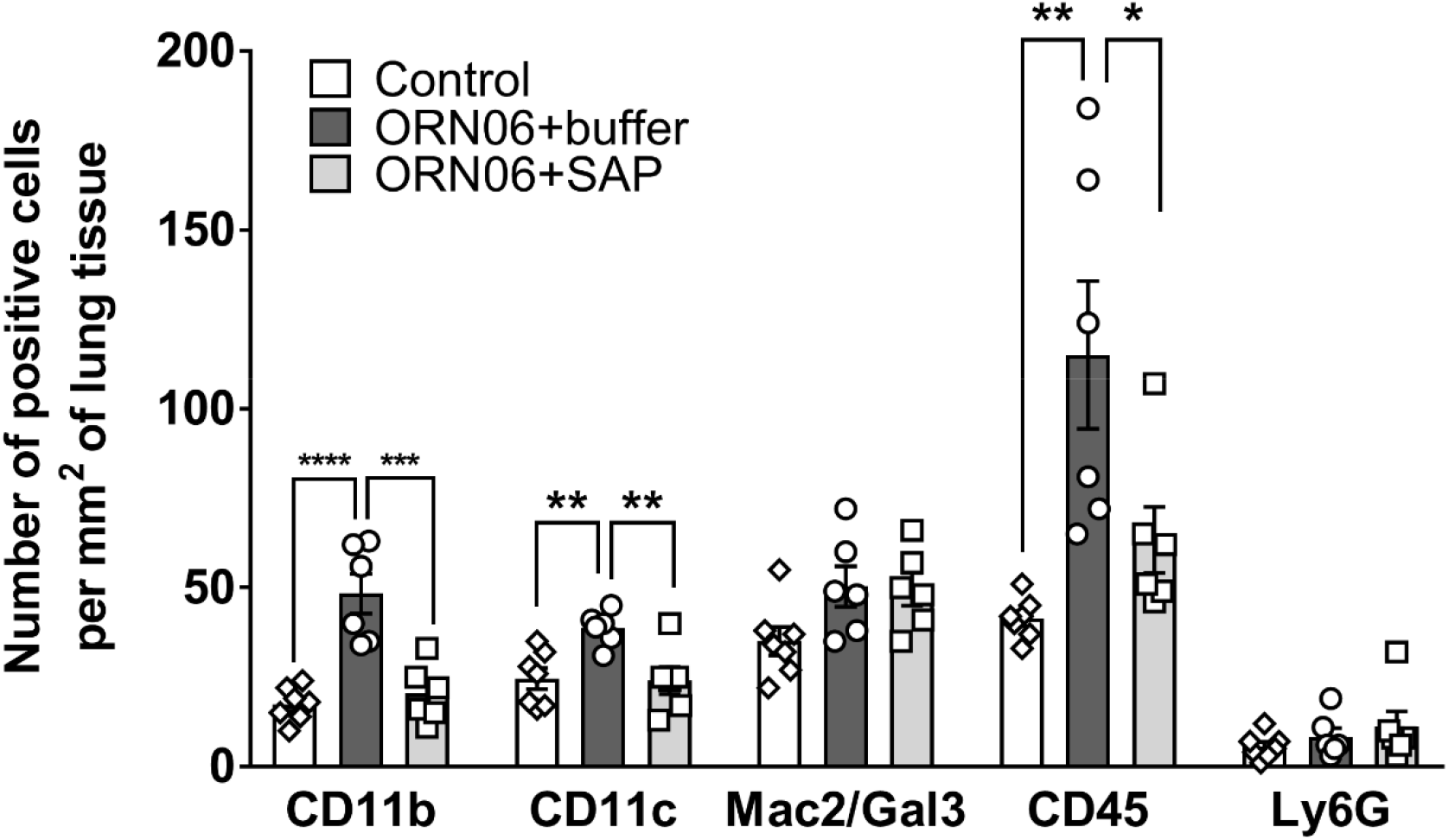
SAP attenuates ORN06-induced upregulation of inflammatory cells in lung tissues post-BAL. Day 3 lung cryosections were stained for the indicated markers. The number of cells stained was determined in 6 randomly chosen 450 µm diameter fields, and the number of positive cells per mm^2^ of lung tissue was calculated for each marker. Values are mean ± SEM, n = 6 mice (3 males and 3 females), except control group where n = 7 (3 males and 4 females). * p < 0.05, ** p < 0.01, *** p < 0.001, and **** p < 0.0001 (1-way ANOVA, Dunnett’s test).

### Aspirated ORN06 increases clot-like aggregates in male mouse BALF, and SAP injections may counteract this

We noticed that the ORN06 treatment with subsequent buffer injections caused the formation of irregular clot-like aggregates visible as dark blue irregular objects in the Wright-Giemsa-stained BALF from 2 of 3 male mice (Fig 4). Defining a clot-like aggregate as an irregularly shaped 2–10 µm object in the BALF staining dark blue with Wright-Giemsa stain, no clot-like aggregates were observed in the BALF cell spots from the 3 male and 5 female control mice or the 3 male and 3 female ORN06-treated with subsequent SAP injections mice. These results indicate that SAP injections may inhibit the formation of clot-like aggregates in the lungs.

**Fig 4.**
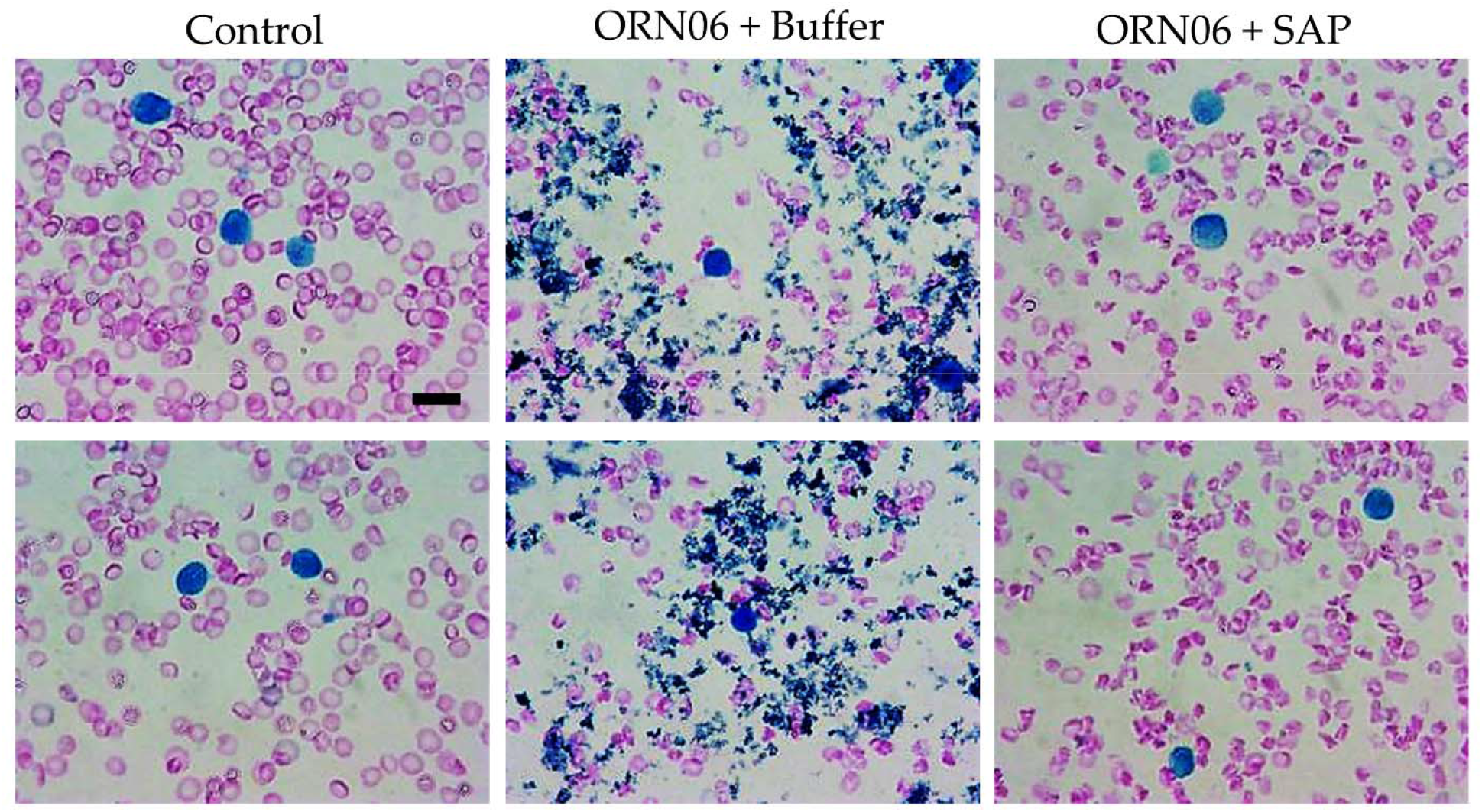
SAP attenuates ORN06-induced clot formation in mouse BAL. Cell spots of BALF at day 3 were stained with Wright-Giemsa stain. The left two images are of BALF cell spots from two different control male mice. The middle two images are of BALF cell spots from two different male mice treated with buffer after ORN06 insult. The right two images are of BALF cell spots from two different male mice treated with SAP after ORN06 insult. Bar is 10 µm.

### Aspirated ORN06 increases alveolar wall thickness and exudates in the alveoli in mice, and SAP injections can counteract this

Compared to the control group, for male and female mice, ORN06 aspirated and buffer-injected mice showed an increase in alveolar wall thickness, and this was reduced by treatment with SAP (Fig 5 A-D and S7 Fig). ORN06 treatment with or without SAP caused a larger increase on alveolar wall thickness in female compared to male mice (S7 Fig). We also noticed that for male and female mice, ORN06 caused the formation of exudates visible as light pink objects in the interior of alveoli (the alveolar airspace) in cryosections stained with hematoxylin & eosin (red arrow, Fig 5B, Fig 5 E and F, and S7 Fig). This effect was reversed by SAP injections (Fig 5A-C, E, and F, and S7 Fig). Compared to female mice, male mice had a greater percentage of alveoli containing ORN06-induced exudates (S7 Fig). Together, these results indicate that ORN06 inhalation increases alveolar wall thickness and the formation of exudates in alveoli, and that SAP injections can inhibit both of these ORN06-induced effects.

**Fig 5.**
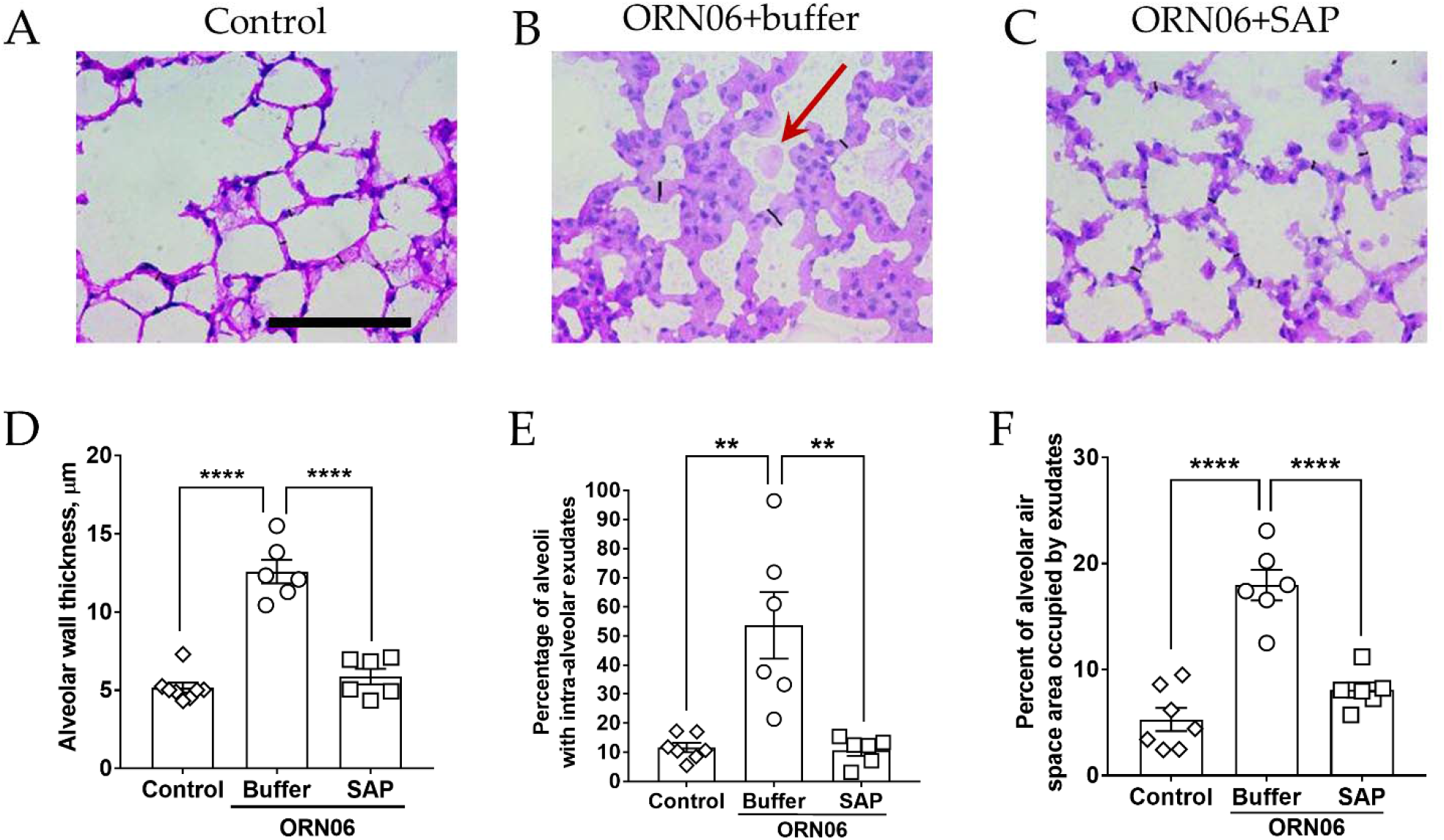
SAP attenuates ORN06-induced alveolar wall thickness. Day 3 lung cryosections of **(A)** control, **(B)** ORN06+buffer, and **(C)** ORN06+SAP were stained with H&E. Images are representative of mice from each group. The thin black lines on the images are the representation of the measured alveolar wall thickness. Scale bar (thick black bar) is 100 µm. Red arrow indicates an exudate in the alveolar air space. **(D)** Quantification of alveolar wall thickness in lung cryosections. For (D), values are mean ± SEM, n = 6 mice (3 males and 3 females) except the control group where n = 8 (3 males and 5 females). **(E)** Quantification of the percentage of alveoli containing one or more exudates in the alveolar airspace observed in lung-section micrographs. **(F)** Quantification of the percentage area of alveoli airspace containing exudate observed in micrographs of lung sections. For (E) and (F), values are mean ± SEM, n = 6 mice (3 males and 3 females) except the control group where n = 7 (3 males and 4 females). For (D), (E), and (F) ** p < 0.01 and **** p < 0.0001 (1-way ANOVA, Bonferroni’s test).

### Aspirated ORN06 increases some serum cytokines in mice, and SAP injections can counteract this

Compared to control, ORN06 treatment of mice caused serum concentrations of IL-1β, IL-6, IL-12p70, IL-23, and IL-27 to increase at day 3, and SAP injections decreased all of these ORN06-induced increases (Fig 6 A-E). Male mice had a somewhat greater ORN06-induced increase in IL-6 than female mice, but did not show an ORN06-induced increase in IL-23 (S8 Fig). ORN06 decreased IL-17 levels, and had no significant effect on levels of several other cytokines, while SAP increased IL-10 and restored IL-17A in ORN06-treated female mice, and decreased MCP-1 in ORN06-treated male mice (S9 and S10 Fig). Together, these results suggest that ORN06 inhalation increases some serum cytokines in mice, and that SAP injections can inhibit these ORN06-induced effects.

**Fig 6.**
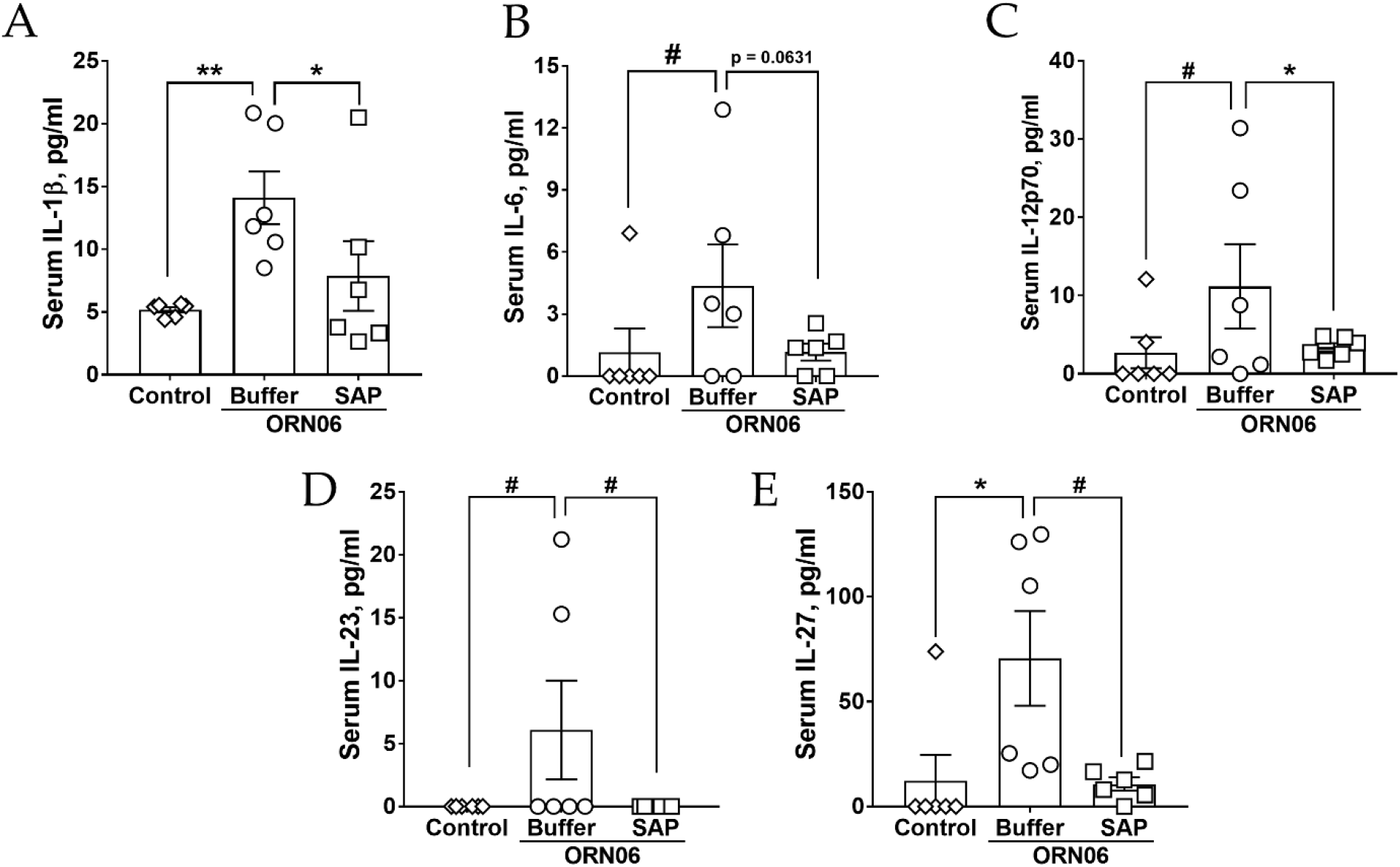
SAP attenuates ORN06-increased serum cytokine levels. Quantification of **(A)** IL-1β, **(B)** IL-6, **(C)** IL-12p70, **(D)** IL-23, and **(E)** IL-27 levels detected in mouse serum for control, ORN06 and then buffer, and ORN06 and then SAP-treated mice. Values are mean ± SEM, n = 6 mice (3 males and 3 females). * p < 0.05 and ** p < 0.01 (t-test). # p < 0.05 (Welch’s t-test).

## Discussion

In this report, we observed that aspiration of ORN06 induced the accumulation of immune cells, alveolar wall thickening, and presence of lung exudates, as previously described when mice were treated with GU-rich ssRNA by tail vein injection (51). Aspiration of ORN06 also led to the formation of clot-like aggregates in some male mouse lung fluids. ORN06 aspiration increased levels of CD11b+, CD11c+, and CD45+ macrophages, CD3-lymphocytes, but not Ly6G positive neutrophils in the lungs, and upregulated IL-1β, IL-6, IL-12p70, IL-23, and IL-27 in the serum. We also observed differences in the response to ORN06 between male and female mice. Unlike tail vein injection, aspiration of the GU-rich ssRNA did not induce death of the mice over 72 hours (51). In mice treated with aspirated ORN06, injections of SAP attenuated lung inflammation, alveolar wall thickening, the presence of lung exudates and clot-like aggregates, and elevated serum cytokines induced by ORN06 aspiration. These results suggest that ORN06 aspiration mimics some of the characteristics of COVID-19, and that SAP could be a potential therapeutic for severe COVID-19 disease and other conditions where a cytokine storm is a clinical problem.

Compared to both healthy controls and patients with viral or bacterial pneumonia, COVID-19 patients, especially severe cases, have in their peripheral blood reduced numbers of CD14+CD16-DR+ “classical” monocytes, which may be due to the increased levels of IL-6 and/or dysregulated myelopoiesis (1, 66-71). In the BAL, patients with severe COVID-19 have more macrophages than mild cases or healthy controls, and the increased macrophage numbers appear to be due to the presence of recruited monocyte-derived inflammatory macrophages (17, 72-74). We found that ORN06-induced increases in BALF cell counts were mirrored by an increase in CD11c+ and CD45+ cells with the morphology of macrophages, suggesting that ORN06 stimulates the accumulation of CD11c+CD45+ alveolar macrophages, but whether this is due to the proliferation of resident alveolar macrophages, migration of CD11c+CD45+ interstitial macrophages into the alveoli, or a rapid phenotypic conversion of recruited CD11b+CD11c- to CD11b-CD11c+ macrophages is unknown (75, 76). Aspiration of ORN06 led to increased numbers of CD11b+, CD11c+, and CD45+ cells, but not Mac2/Gal3 positive cells, retained in the lungs after BAL. As Mac2/Gal3 is a marker for tissue resident lung macrophages (65), these data suggest that ORN06 generates a response that induces the recruitment of CD11b+CD45+ inflammatory macrophages, which may undergo phenotypic conversion to CD11b-CD11c+CD45+ macrophages, and/or ORN06 induces the proliferation of lung resident CD11c+CD45+ interstitial macrophages (75-77). Injections of SAP attenuated the accumulation of macrophages in both the BAL fluid and in the post-BAL lung tissue. These data suggest that SAP can be used to target the recruited macrophage populations without altering the tissue resident macrophages, or elevating the neutrophils in the lung.

ORN06 aspiration also led to a small but significant increase in the number of BALF lymphocytes, which were CD3 negative, suggesting this was due to either the recruitment of CD3 negative natural killer (NK) cells or other types of innate lymphoid cells (ILC), or the proliferation of lung resident ILCs (78-80). SAP was also effective at attenuating the ORN06-induced increase in BALF lymphocytes.

COVID-19 patients have reduced numbers of ILC, including NK cells, in the peripheral blood, but scRNA-seq analysis from COVID-19 patients indicate higher numbers of NK cells in the BAL (70). NK cells and other ILCs are potent producers of a variety of cytokines, depending on the subtype of ILCs, stimulus, and tissue where the ILCs are found, but how the changes in the number and composition of ILCs subsets affect cytokine levels in COVID-19 is still unclear (1, 81). COVID-19 is also characterized by a “cytokine storm” with increased levels of pro-inflammatory cytokines in the blood, such as IL-1β, TNF-α, IL-6, IL-12 (67, 70, 71, 82), and we detected increased IL-1β, IL-6, IL-12, IL-23, and IL-27 in the serum following ORN06 aspiration. Injections of SAP attenuated the increased levels of the above cytokines, suggesting that SAP can either induce cells to degrade extracellular cytokines, or inhibit cells from secreting cytokines the above cytokines. SAP also elevated the serum levels of the “anti-inflammatory cytokine” IL-10, as observed previously (27, 35, 41). Previous studies suggest that IL-17A plays an important role in enhancing antiviral T helper type 1 responses in the female genital tract, where female mice lacking IL-17A have lower proportions of IFN-γ+ T_h_1 cells than wild type female mice (83). Compared to male mice, female mouse bladders have higher IL-17 expression (84).We observed that ORN06 decreased serum IL-17A in female but not male mice, and that SAP reversed the ORN06 effect in female mice. Together, this suggests that SAP can reverse the effects of ORN06 on several serum cytokines.

Severe COVID-19 disease is 1.5 to 2 times higher in men than women (4-8).We found that ORN06 aspiration led to a higher BALF cell numbers, clot-like aggregates, and alveoli containing exudates in male mice compared to female mice, which suggests a similar response in mice compared to humans. Many diseases are associated with sex-based differences in disease incidence, prevalence, and severity (85-87), but the causes of these differences for COVID-19 in particular remain unclear.

Together, our results and the work described above suggest the intriguing possibility that aspiration of oligonucleotides such as ORN06 could be a simple model for COVID-19 infection. In addition, SAP strongly attenuated the effects of aspiration of ORN06, suggesting that SAP could be a potential therapeutic for severe COVID-19 disease and possibly other diseases where a cytokine storm is a contributing factor for disease severity or progression.

## Supplementary figures and supplementary figure legends

**S1 Fig.**
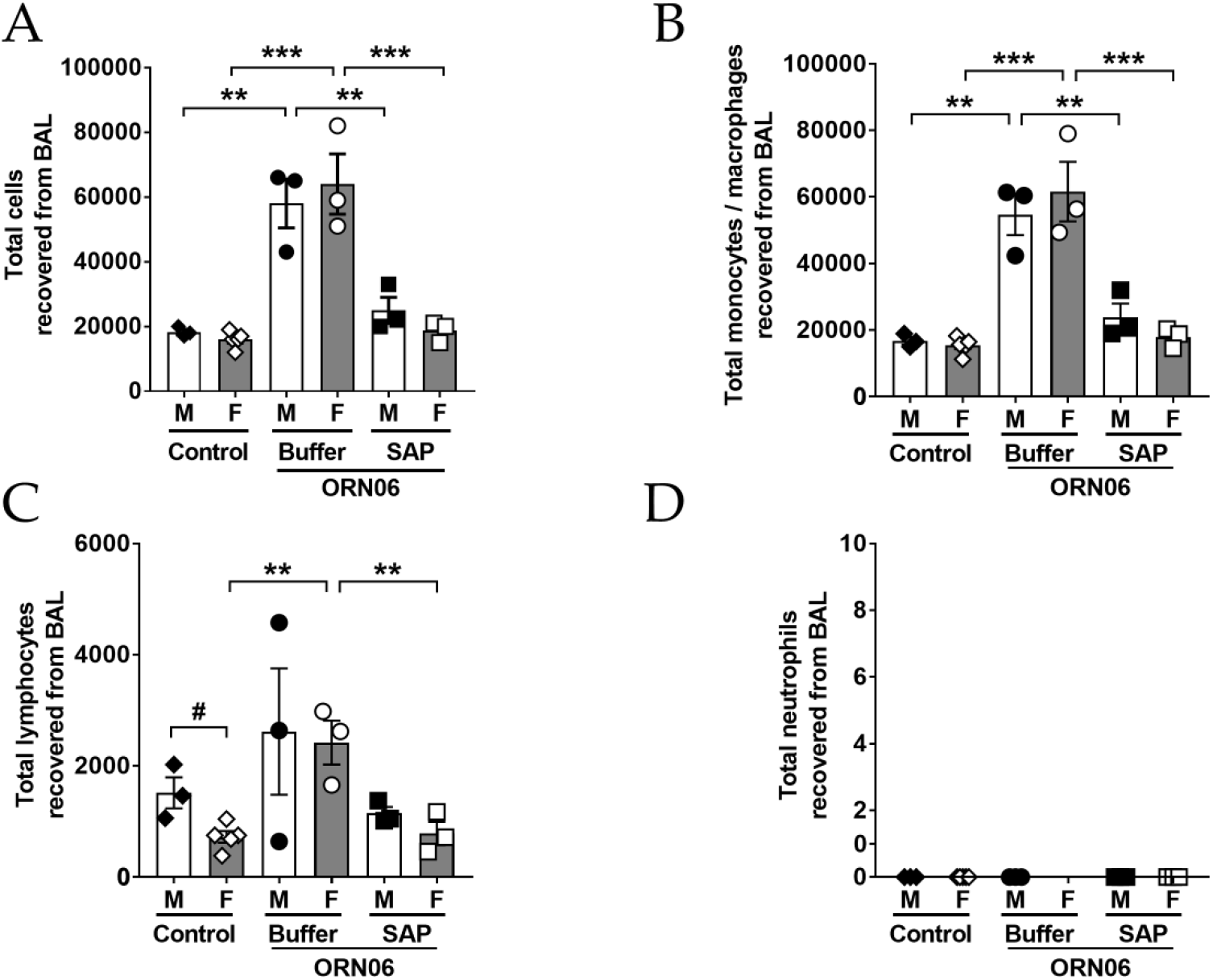
Cells in male and female mouse BAL. The data from Figure 1A-D were separated for male (M) and female (F) mice from each group for **(A)** total cells, **(B)** monocytes/ macrophages, **(C)** lymphocytes, and **(D)** neutrophils. Values are mean ± SEM. For male mice n = 3 in each group and for female mice n = 3 except for female mice control group, where n = 5. ** p < 0.01 and *** p < 0.001 (1-way ANOVA, Bonferroni’s test). # p < 0.05 (t-test).

**S2 Fig.**
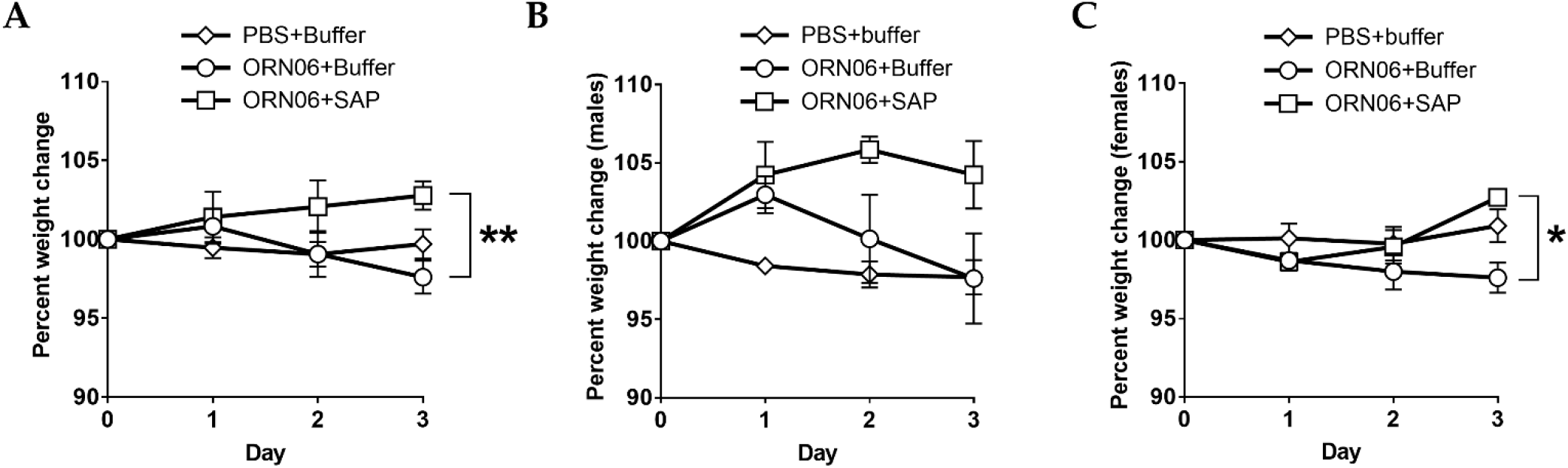
SAP treated mice are resistant to a decline in body weight after ORN06 aspiration. Percent change in body weight for **(A)** both males and females, **(B)** males, and **(C)** females after the indicated treatments. Values are mean ± SEM, n = 6 (3 males and 3 females) except for PBS-aspirated and then PBS-treated, where n=8 (3 males and 5 females). * p < 0.05 and ** p < 0.01 (1-way ANOVA, Bonferroni’s test).

**S3 Fig.**
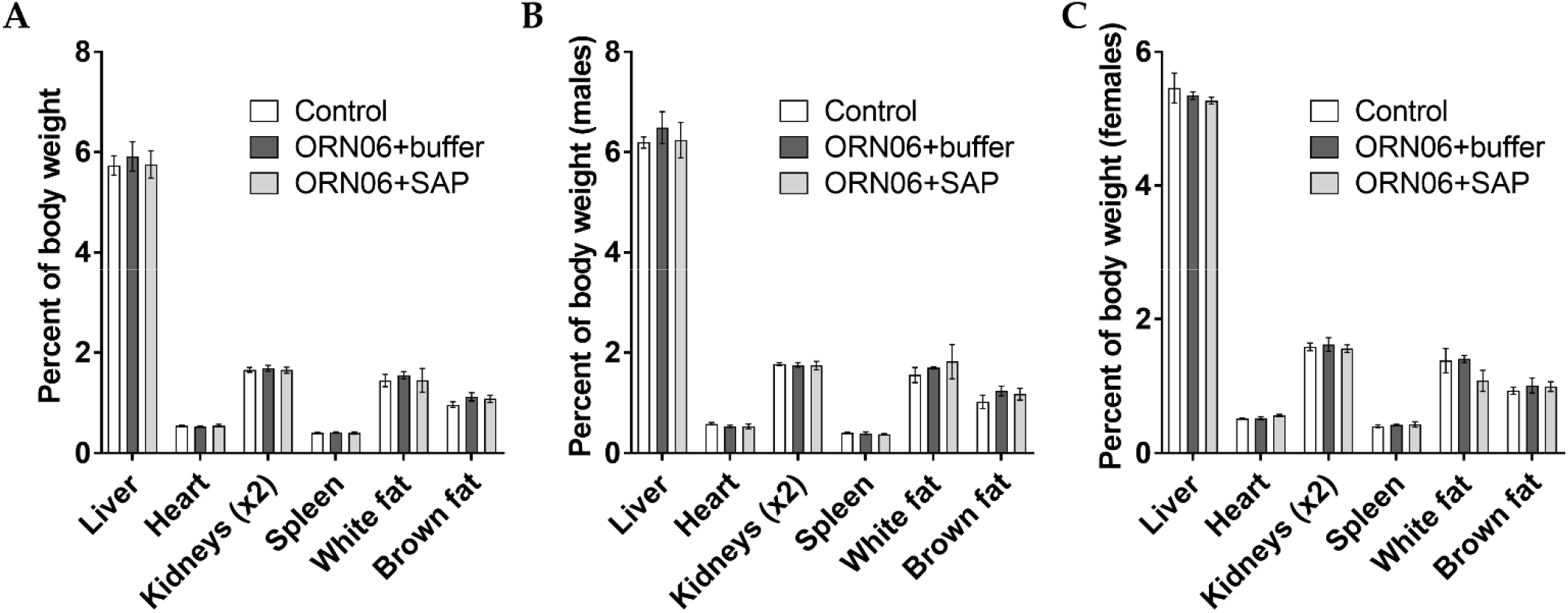
Organ weights. Organ weights as percent of total body weight at day 3 of **(A)** both male and female mice, **(B)** males, and **(C)** females after the indicated treatments. Values are mean ± SEM, n = 6 (3 males and 3 females) except for control, where n = 8 (3 males and 5 females).

**S4 Fig.**
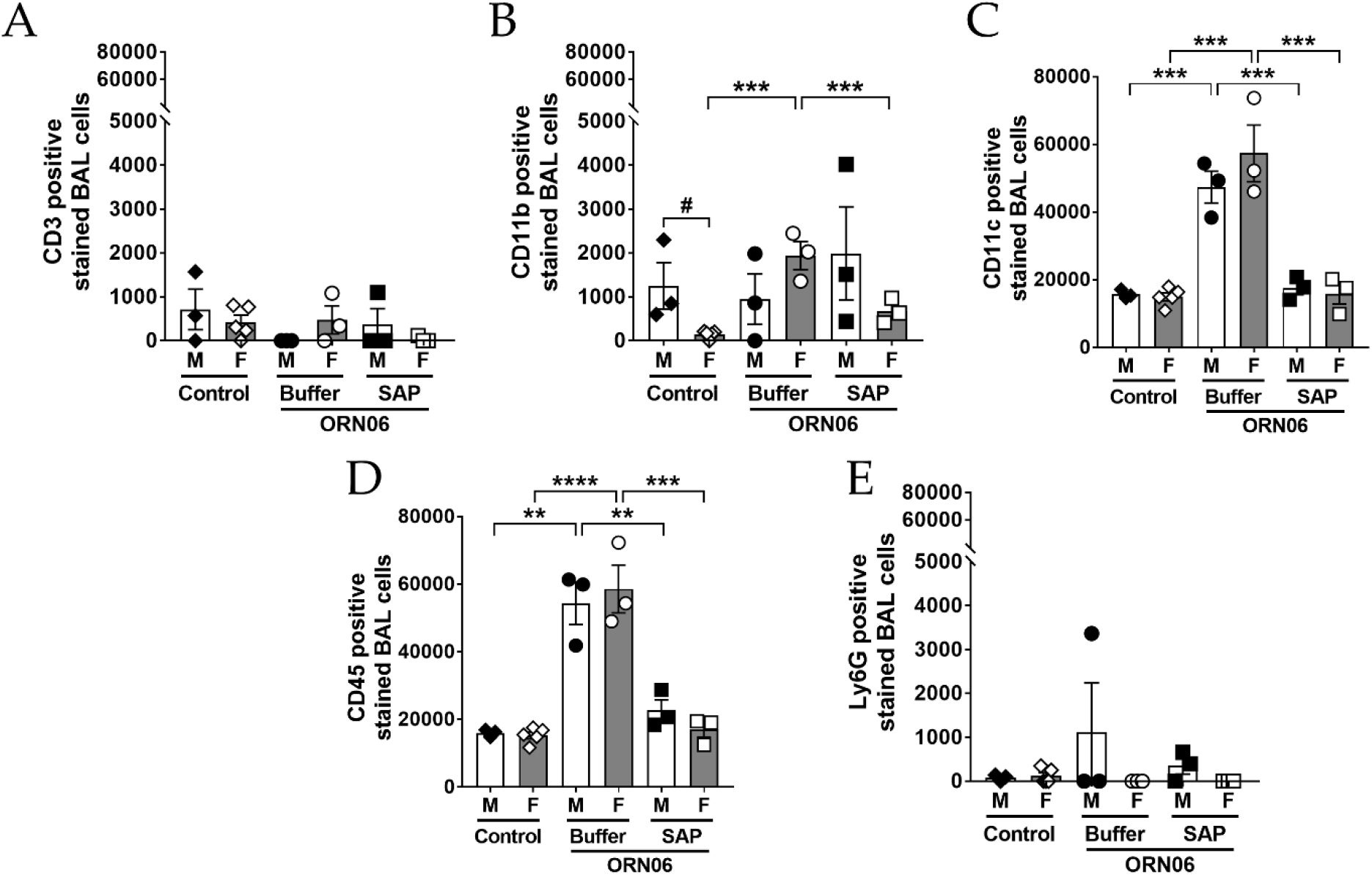
Specific cell types in male and female mouse BAL. The data from Figure 2 was separated for **(A)** CD3 positively stained cells, **(B)** CD11b positively stained cells, **(C)** CD11c positively stained cells, **(D)** CD45 positively stained cells, and **(E)** Ly6G positively stained cells from male (M) and female (F) mice from each group. Values are mean ± SEM. For male mice n = 3 and for female mice n = 3 except for female mice control group, where n = 5. ** p < 0.01, *** p < 0.001, and **** p < 0.0001 (1-way ANOVA, Bonferroni’s test). # p < 0.05 (t-test).

**S5 Fig.**
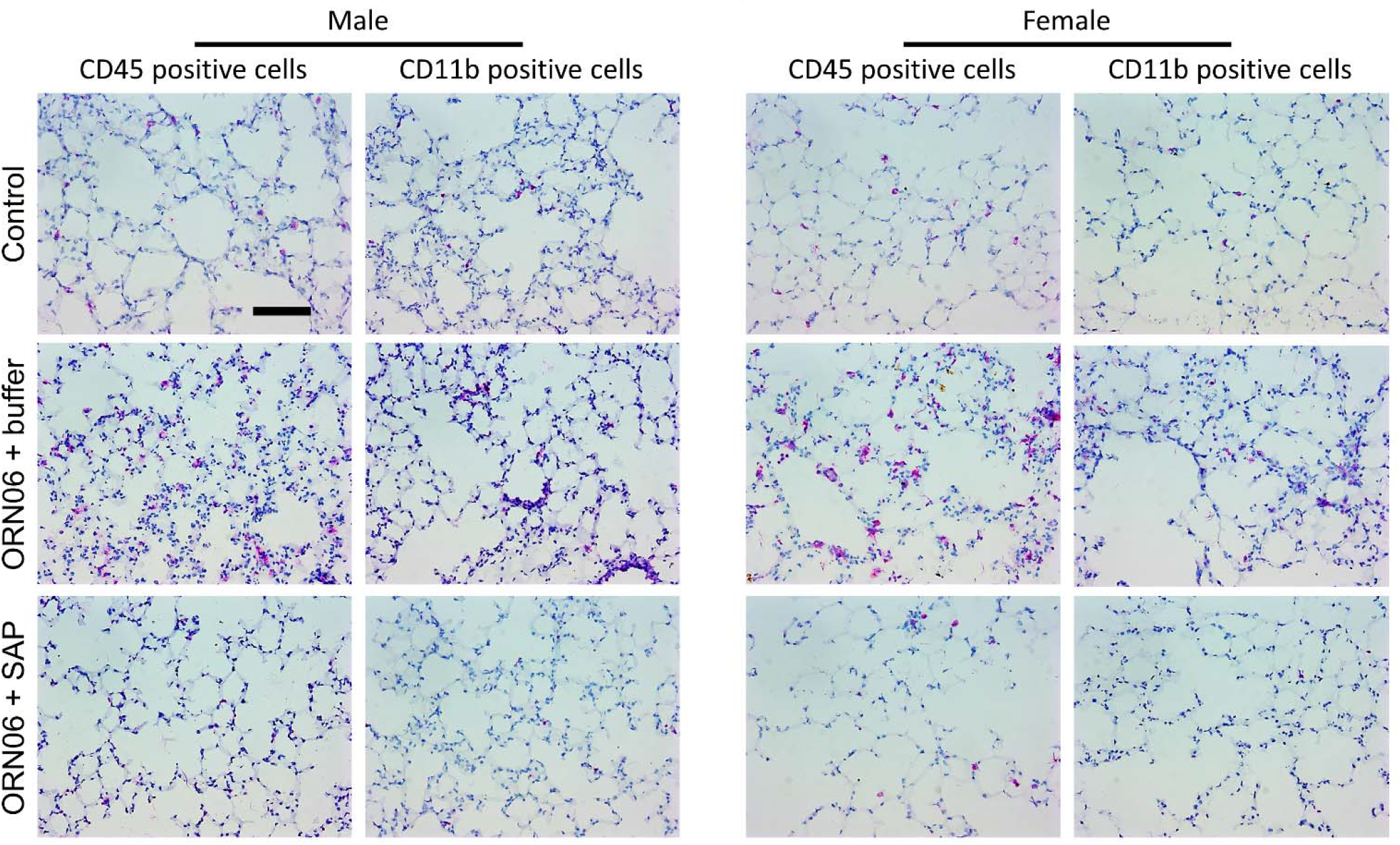
Images of stained post-BAL lung cryosections. Representative images of male and female mouse lung cryosections stained with antibodies against CD45 and CD11b. Red is staining, blue is counter stain. Bar is 100 μm. Images are representative of n = 3 in each male and female group, except for female mouse control group, where n = 4.

**S6 Fig.**
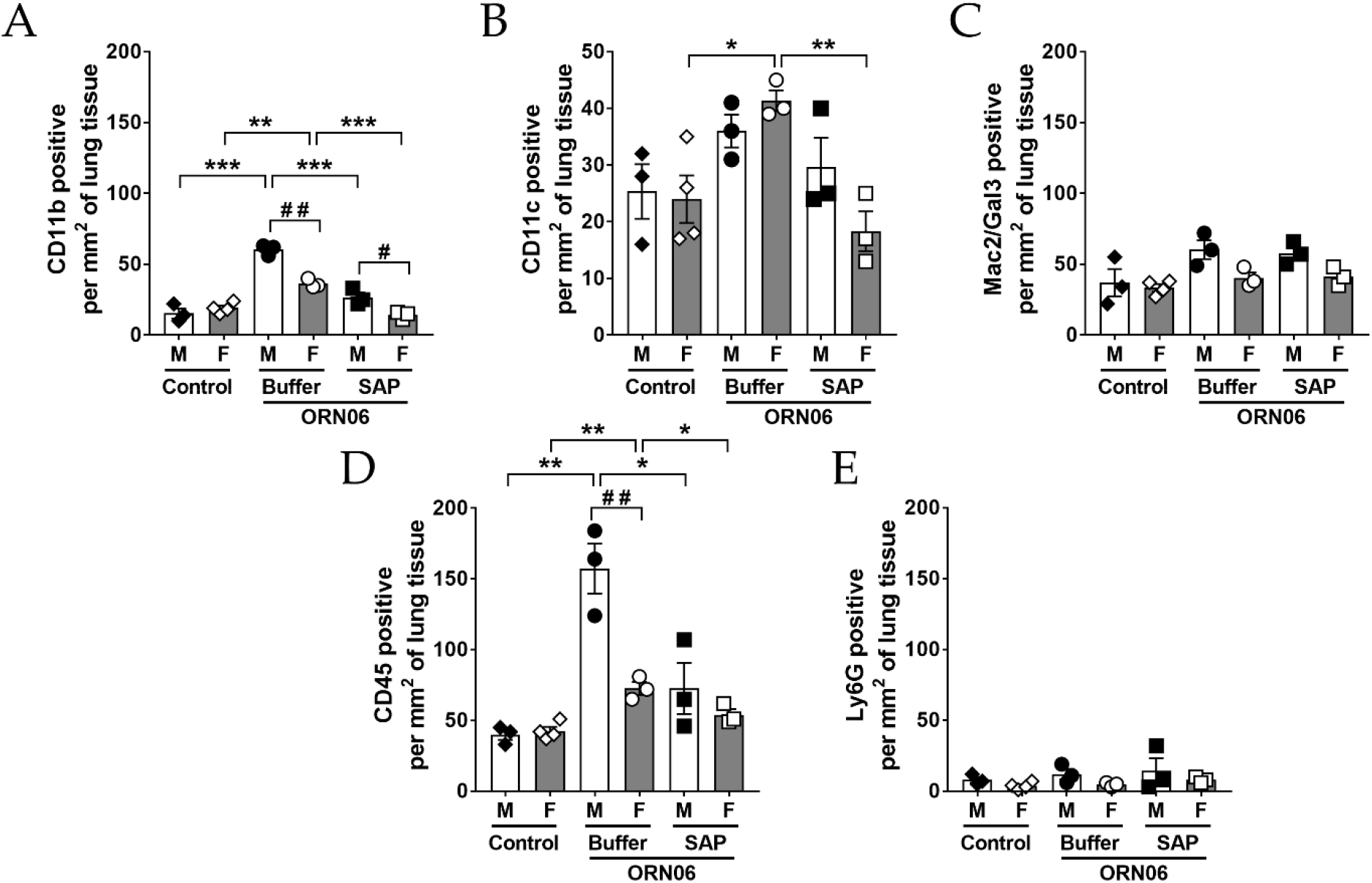
Specific cell types in male and female mouse post-BAL lung cryosections. The data from Figure 3 was separated for **(A)** CD11b positively stained cells, **(B)** CD11c positively stained cells, **(C)** Mac2/Gal3 positively stained cells, **(D)** CD45 positively stained cells, and **(E)** Ly6G positively stained cells from male (M) and female (F) mice from each group. Values are mean ± SEM. For male mice n = 3 and for female mice n = 3 except for female mice control group, where n = 4. * p < 0.05, ** p < 0.01, and *** p < 0.001 (1-way ANOVA, Dunnett’s test). # p < 0.05 and ## p < 0.01 (t-test).

**S7 Fig.**
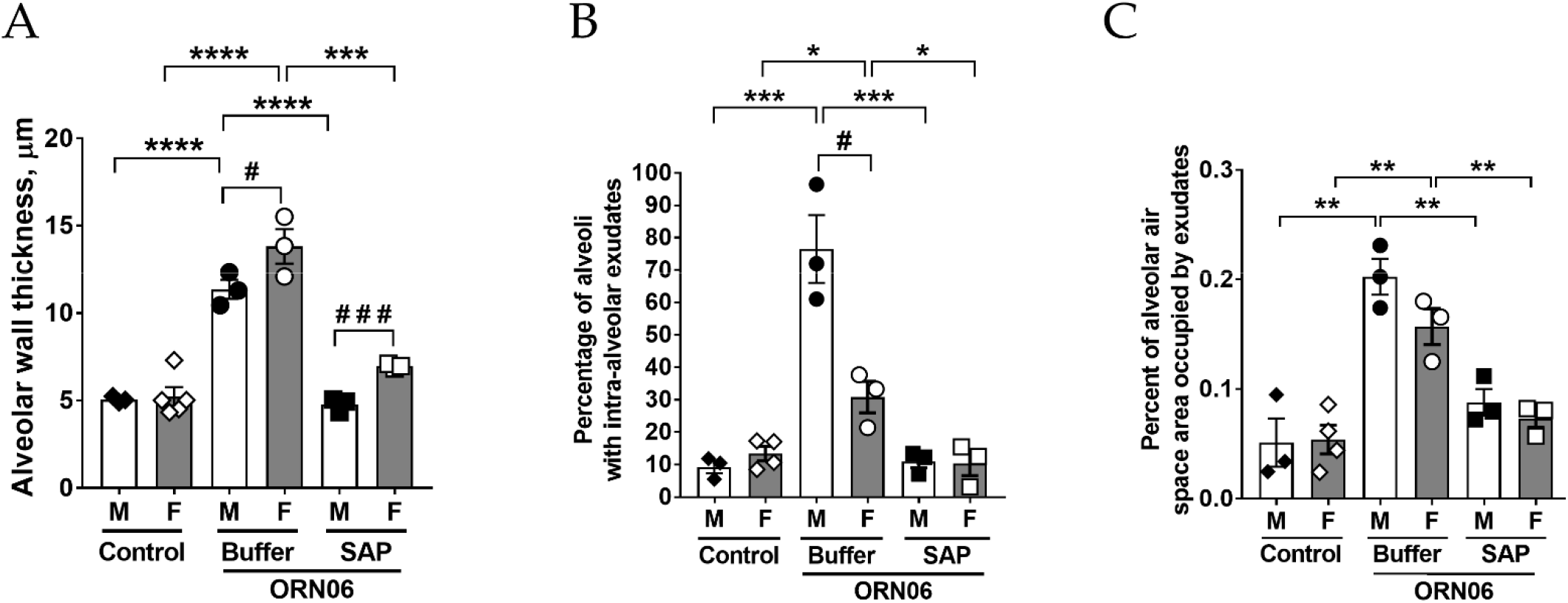
Alveolar wall thickness and alveolar exudates in in male and female mice. The data from Figures 5 D, E and F were separated for **(A)** alveolar wall thickness, **(B)** percent of alveoli with intra-alveolar exudates, and **(C)** percent of alveolar airspace area occupied by exudates from male (M) and female (F) mice from each group. Values are mean ± SEM. For male mice n = 3 in each group and for female mice n = 3 except for female mice control group in (A), where n = 5 and in (B – C), where n = 4. * p < 0.05, ** p < 0.01, *** p < 0.001, and **** p < 0.0001 (1-way ANOVA, Bonferroni’s test). # p < 0.05, ### p < .001 (t-test).

**S8 Fig.**
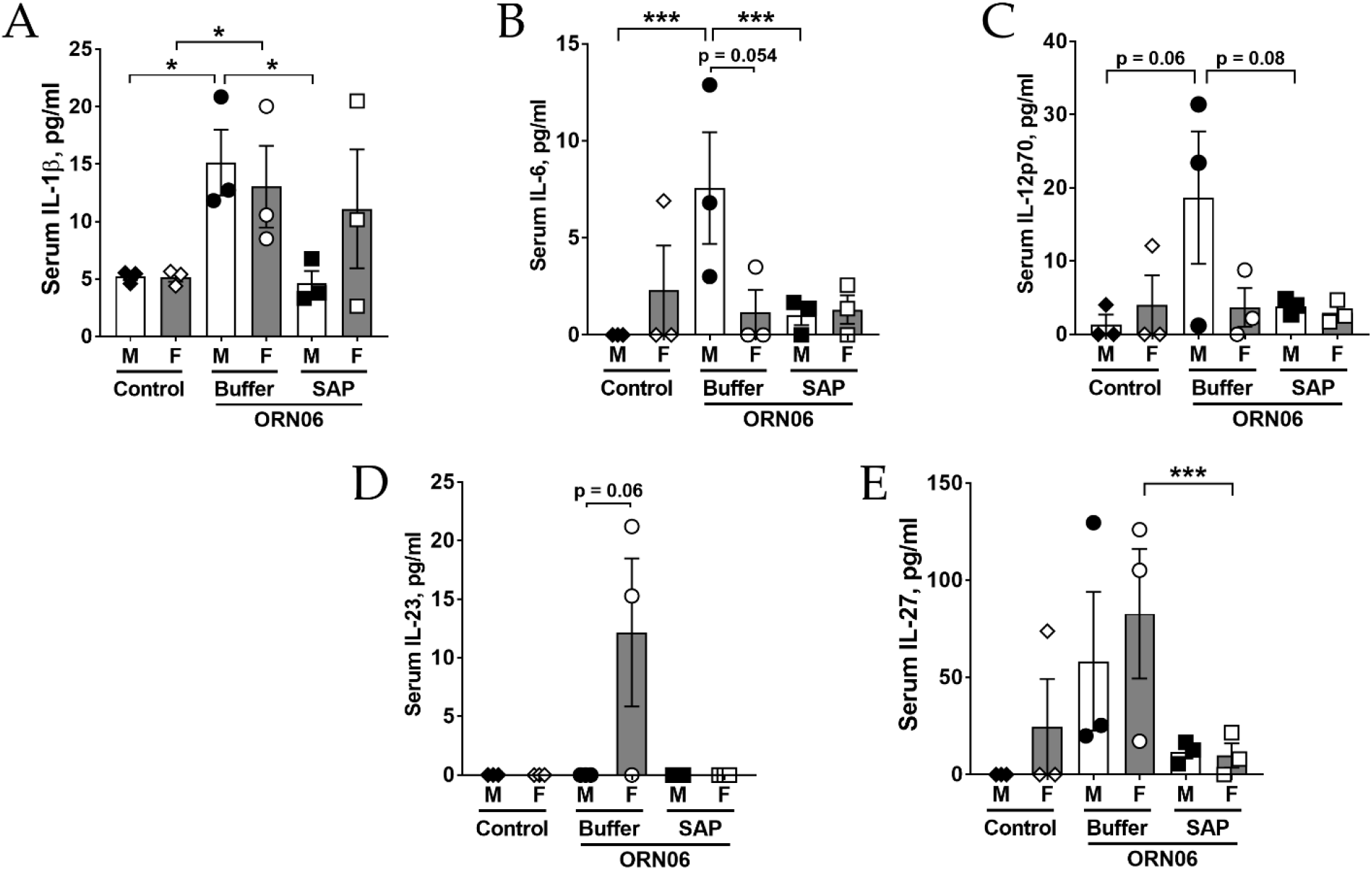
Cytokine levels from male and female mouse sera. The data from Figure 6 A, B, C, D, and E were separated for serum **(A)** IL-1β, **(B)** IL-6 **(C)** IL-12p70, **(D)** IL-23, and **(E)** IL-27 levels from male (M) and female (F) mice from each group. Values are mean ± SEM. For male mice and female mice n = 3 in each group. * p < 0.05 and *** p < 0.001 (t-test).

**S9 Fig.**
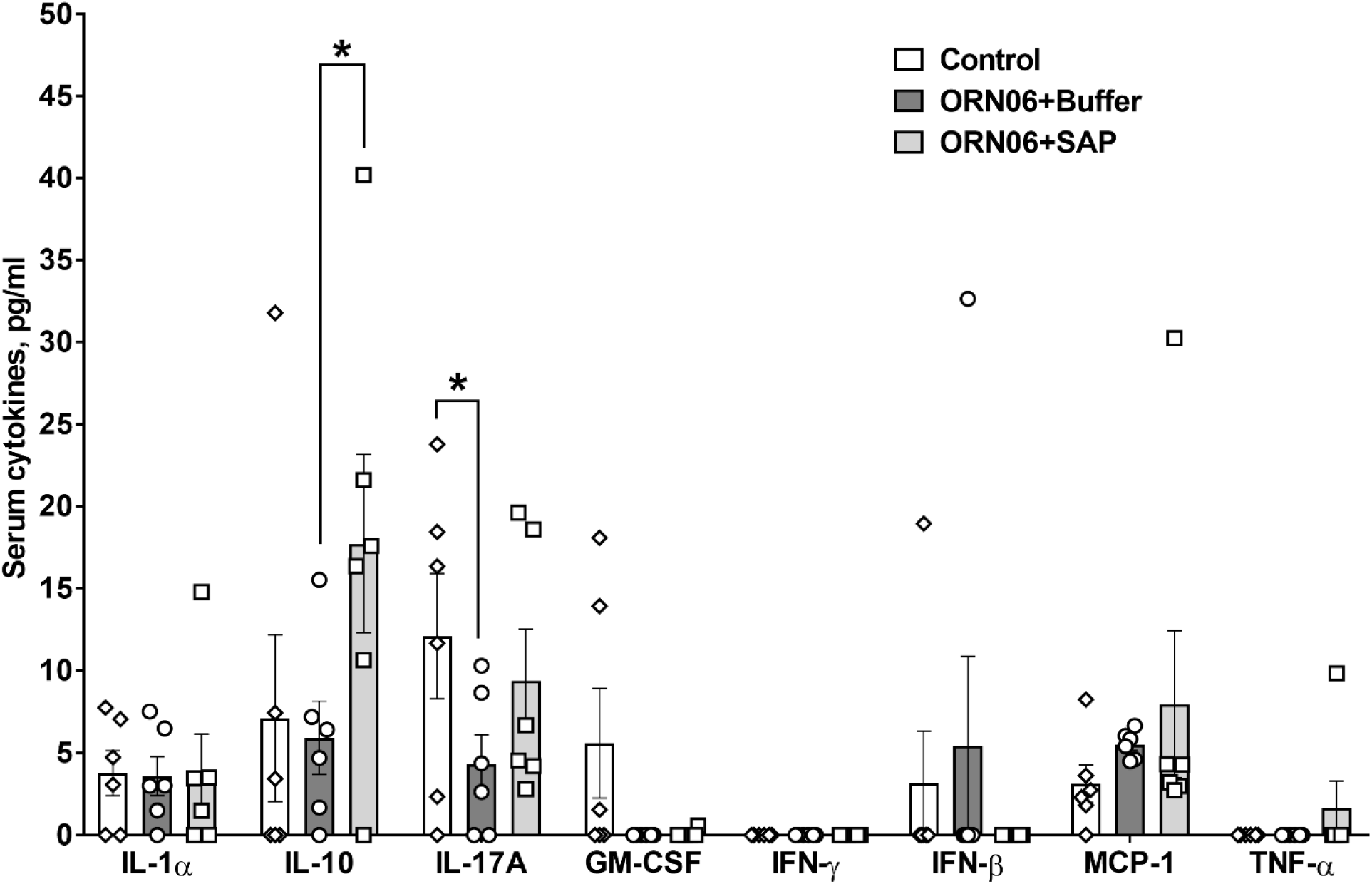
Cytokine levels from mouse sera. Sera were collected at day 3 and assayed by ELISA for the indicated cytokines. Values are mean ± SEM, n = 6 (3 males and 3 females). * p < 0.05 (1-way ANOVA, Bonferroni’s test).

**S10 Fig.**
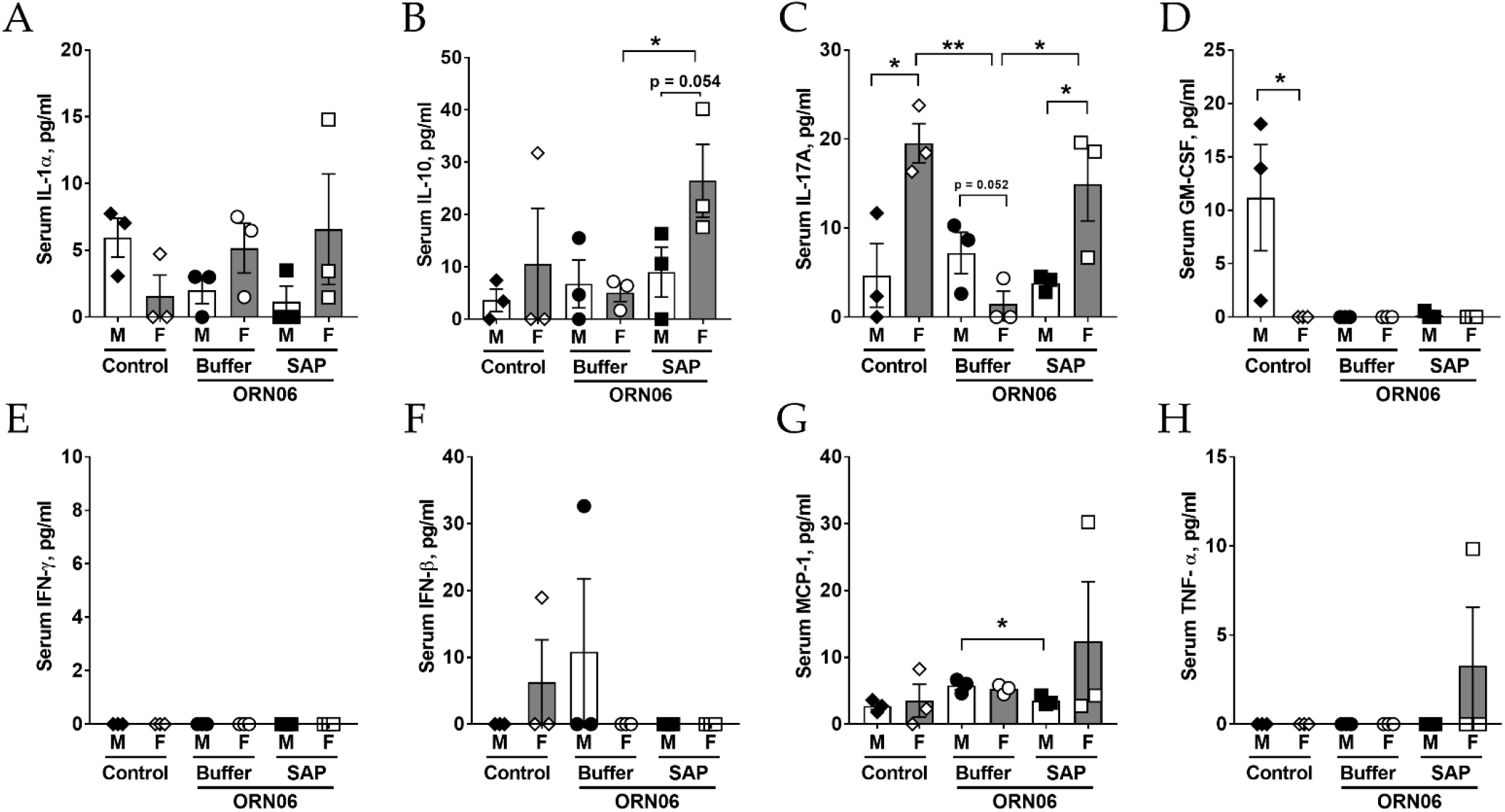
Cytokine levels from male and female mice sera. The data from Supplementary figure 9 were separated for serum **(A)** IL-1α, **(B)** IL-10 **(C)** IL-17, **(D)** GM-CSF, **(E)** IFN-γ, **(F)** IFN-β, **(G)** MCP-1, and **(H)** TNF-α levels from male (M) and female (F) mice from each group. Values are mean ± SEM. For male mice and female mice n = 3 in each group. * p < 0.05 and ** p < 0.01 (t-test).

## Notes

**Conflict of Interest statement:** Texas A&M University has a patent application on the use of SAP to treat cytokine storm and COVID-19-associated lung inflammation and lung damage. DP, TRK, and RHG are inventors on this patent application.

Funding statement: This work was supported by NIH HL-132919 and NIH GM118355-03S1

### Competing Interest Statement

Texas A&M University has a patent application on the use of SAP to treat cytokine storm and COVID-19-associated lung inflammation and lung damage.
DP, TRK, and RHG are inventors on this patent application.

